# Induction of the nicotinamide riboside kinase NAD^+^ salvage pathway in skeletal myopathy of H6PDH KO mice

**DOI:** 10.1101/567297

**Authors:** Craig L. Doig, Agnieszka E. Zielinska, Rachel S. Fletcher, Lucy A. Oakey, Yasir S. Elhassan, Antje Garten, David Cartwright, Silke Heising, Ahmed Alsheri, David G. Watson, Jerzy Adamski, Daniel A. Tennant, Gareth G. Lavery

## Abstract

**Background:** Hexose-6-Phosphate Dehydrogenase (H6PDH) is a generator of NADPH in the Endoplasmic/Sarcoplasmic Reticulum (ER/SR). Interaction of H6PDH with 11β-hydroxysteroid dehydrogenase type 1 provides NADPH to support oxo-reduction of inactive to active glucocorticoids, but the wider understanding of H6PDH in ER/SR NAD(P)(H) homeostasis is incomplete. Muscle specific lack of H6PDH results in a deteriorating skeletal myopathy, altered glucose homeostasis, ER stress and activation of the unfolded protein response. Here we further assess muscle responses to H6PDH deficiency to delineate pathways that may underpin myopathy and link SR redox status to muscle wide metabolic adaptation.

**Methods:** We analysed skeletal muscle from H6PDH knockout (H6PDKO), H6PDH and NRK2 double knockout (DKO) and wild-type (WT) mice. H6PDKO mice were supplemented with the NAD^+^ precursor nicotinamide riboside. Skeletal muscle samples were subject to biochemical analysis including NAD(H) measurement, LC/MS based metabolomics, Western immunoblotting, and high resolution mitochondrial respirometry. Genetic and supplement models were assessed for degree of myopathy compared to H6PDKO.

**Results:** H6PDKO skeletal muscle showed adaptations in the routes regulating nicotinamide and NAD^+^ biosynthesis, with significant activation of the Nicotinamide Riboside Kinase 2 (NRK2) pathway. Associated with changes in NAD^+^ biosynthesis H6PDKO muscle had impaired mitochondrial respiratory capacity with altered mitochondrial acylcarnitine and acetyl CoA metabolism. Boosting NAD^+^ levels through the NRK2 pathway using the precursor nicotinamide riboside had no effect to mitigate ER stress and dysfunctional mitochondrial respiratory capacity or Acetyl-CoA metabolism. Similarly, H6PDKO/NRK2 double KO mice did not display an exaggerated timing or severity of myopathy or overt change in mitochondrial metabolism despite depression of NAD^+^ availability.

**Conclusions:** These findings suggest a complex metabolic response to changes to muscle SR NADP(H) redox status that result in impaired mitochondrial energy metabolism and activation of cellular NAD^+^ salvage pathways. It is possible that SR can sense and signal perturbation in NAD(P)(H) that cannot be rectified in the absence of H6PDH. Whether NRK2 pathway activation is a direct response to changes in SR NAD(P)(H) availability or adaptation to deficits in metabolic energy availability remains to be resolved.

## Background

Our understanding of NADP(H) redox homeostasis within muscle sarcoplasmic reticulum (SR) and its influence over cellular metabolism is developing^1,2^. Hexose-6-phosphate dehydrogenase (H6PDH) is an SR luminal enzyme that uses glucose-6-phosphate to produce NADPH^3,4^. A known function of H6PDH is its physical interaction with 11β-hydroxysteroid dehydrogenase type 1 (11β-HSD1) in the SR lumen^5^. H6PDH NADPH generation supports 11β-HSD1 oxo-reductase activity for glucocorticoid production^6^. The importance of H6PDH-11β-HSD1 interaction is revealed in humans with ‘apparent’ cortisone reductase deficiency (ACRD) due to inactivating mutations in the *H6PD* gene^7,8^. In ARCD the lack of H6PDH and NADPH switches 11β-HSD1 activity towards glucocorticoid oxidation resulting in increased tissue glucocorticoid clearance and relative insensitivity. Affected individuals manifest disease as a function of increasing hypothalamic-pituitary-adrenal axis activity and resultant excess adrenal androgen production^9^. H6PDH knockout (H6PDKO) mice endorse the biochemistry with manifestation of an ARCD-like phenotype, showing tissue glucocorticoid insensitivity and HPA axis activation^4^. Liver and adipose tissues show phenotypes associated with relative glucocorticoid insensitivity. The liver has impaired ability to conduct gluconeogenesis and with increased glycogen synthesis rates, while adipose tissue has impaired ability to store and mobilise lipid^10,11^.

Skeletal muscle of H6PDH knockout (H6PDKO) mice develop a deteriorating skeletal myopathy associated with large intrafibrillar membranous vacuoles, abnormal sarcoplasmic reticulum (SR) structure, dysregulated expression of SR proteins involved in calcium metabolism and switching of type II to type I fibers and impaired force generation^12^. Associated with myopathy is ER stress and activation of the unfolded protein response-partially attributed to altered SR lumen protein folding capacity^12,13^. Metabolically H6PDKO are hypoglycaemic on fasting, more insulin sensitive with increased insulin-stimulated glucose uptake in type II fiber rich muscle and increased glycogen content^12,14^. Importantly, H6PDKO myopathy is independent of the role of 11β-HSD1 and glucocorticoids^14^.

It is proposed that H6PDH contributes more to ER/SR redox balance beyond regulation of 11β-HSD1 activity as H6PDH metabolism of G6P, and effects on the ER lumen NADPH/NADP+ ratio, can impact glucose production and adrenal steroidogenic enzyme activity^15^. Less clear is the interaction and contribution of H6PDH and ER/SR NADP(H) to whole cell redox and cellular energy metabolism.

Myopathy in H6PDKO mice is seemingly uncoupled from glucocorticoid metabolism and sensitivity. We wanted to investigate primary pathways responding to the perturbations in SR NADP(H) redox due to H6PD deficiency using unbiased and targeted metabolomics and existing transcriptional data^12^. We show that in young mice, prior to overt phenotypic presentation, H6PDKO remodels NAD(P)(H) biosynthesis, particularly through the nicotinamide riboside kinase 2 (NRK2) pathway. H6PDH deficiency also results in upregulation of pathways to acetyl-CoA production leading to accumulation of short chains acylcarnitines, elevated acetylation of mitochondrial proteins and impaired mitochondrial fatty acid oxidation. Acetylation of the mitochondrial proteome is associated with depression in energy production, and can be countered through the activity of the NAD^+^-dependent deacetylase SIRTs^16,17^. Taking advantage of elevated NRK2 expression we augmented NAD^+^ availability in H6PDKO mice by supplementing the precursor nicotinamide riboside (NR). Similarly H6PD/NRK2 double KO mice were examined as a means to further abrogate NAD^+^ availability. While each approach was able to modulate NAD^+^ metabolism, neither was able to rescue or exacerbate mitochondrial dysfunction or myopathy. These findings reveal a complex metabolic response to changes to muscle SR NADP(H) redox status and associated impairment in mitochondrial energy metabolism.

## Materials and Methods

### Animal care, Mouse strains, storage

H6PDH KO mice (C57/BL5J background) were group housed in a standard temperature (22°C) and humidity-controlled environment with 12:12-hour light:dark cycle. Nesting material was provided and mice had ad libitum access to water and standard chow. Mice were sacrificed using schedule one cervical dislocation and tissues were immediately. Collections were all performed at 10-11am.

Intraperitoneal injections of Nicotinamide Riboside (400mg/kg) were given twice daily. (10am and 3pm) for 4 days before cervical dislocation and skeletal muscle bed collection on Day 5 (10am), tissues were flash frozen, fixed or for high resolution respirometry stored in BIOPS buffer. Heterozygotes of H6PDKO and NRK2KO were breed to generate H6PD-NRK2 KO mice. Mice were utilised between 8-12 weeks of age. Experiments were consistent with current UK Home Office regulations in accordance with the UK Animals Scientific Procedures Act 1986.

### RNA extraction and qRT-PCR

Total RNA was extracted from skeletal muscle tissue using TRI-reagent (Invitrogen). RNA quality was determined by visualisation on a 1.5% agarose gel and quantity was measured by nanodrop absorbance at 260nm. Reverse transcription was conducted using 1μg RNA that was incubated with 250uM random hexamers, 5.5mM MgCl_2_, 500uM dNTPs, 20units RNase inhibitor 63units multi-scribe reverse transcriptase and 1x reaction buffer. Reverse transcription was performed on a thermocycler set at the following conditions: 25°C 10minutes, 48°C for 30 minutes before the reaction was terminated by heating to 98°C for 5 minutes. cDNA levels were determined using an ABI7500 system (Applied Biosystems), reactions were conducted in a 384-well plate in single-plex format. Primers and probes were purchased as Assay on Demand (FAM) products (Applied Biosystems). Total reaction volumes used were 10ul containing Taqman Universal PCR mix (Applied Biosystems). All reactions were normalised to 18s rRNA (VIC) (Applied Biosystems). The real-time PCR reaction was performed at the following conditions: 95°C for 10 minutes then 40 cycles of 95°C for 15 seconds and 60°C for 1 minute. Data were collected as Ct values and used to obtain deltaCt (dCt) values and expressed as fold change ±standard error of the mean(SEM).

### Mitochondrial DNA (mtDNA) Copy number

DNA was isolated using phenol-chloroform and washed in Ethanol before quantification using nanodrop. DNA from all samples was diluted to 50ng/μl and subject to real-time PCR using SYBRgreen master mix (Applied Biosystems) using the following primers specific for nuclear and mitochondrial DNA.

Nuclear DNA: TERT Forward CTAGCTCATGTGTCAAGACCCTCTT Reverse GCCAGCACGTTTCTCTCGTT, Mitochondrial DNA: DLOOP Forward AATCTACCATCCTCCGTGAAACC Reverse TCAGTTTAGCTACCCCCAAGTTTAA, ND4 Forward AACGGATCCACAGCCGTA Reverse AGTCCTCGGGCCATGATT.

### Western Immunoblotting

Protein lysates were collected in RIPA buffer (50mmol/l Tris pH 7.4, 1% NP40, 0.25% sodium deoxycholate, 150mmol/l NaCl, 1mmol/l EDTA), protease & phosphatase inhibitor cocktail (Roche, Lewes, U.K.), stored at −80°C (>30 minutes), defrosted on ice and centrifuged at 4°C (10mins, 12000 rpm). The supernatant was recovered and total protein concentration was assessed by Bio-Rad assay. Total proteins were resolved on a 12% SDS-PAGE gel and transferred onto a nitrocellulose membrane. Primary antibodies; Anti-Goat H6PDH (in house), Anti-Mouse NAMPT (Bethyl A300-372A), Anti-Rabbit NRK1 (in house), Anti-Rabbit NRK2 (in house), Anti-Rabbit CHOP (Cell Signaling D46F1 5554), Total Lysine Ac (Cell Signaling 9441), Anti-Mouse Rodent OXPHOS Mitoprofile (Abcam ab110413), IDH2 K413ac (GeneTel AC0004), IDH2 Total (Abcam ab55271), H3K9ac (Cell Signaling 9649), H3K18ac (Cell Signaling 9675), ERO1a (Cell Signaling 3264), PDI (Cell Signaling 3501), alpha-Tubulin (Santa Cruz B-7 sc-5286). Secondary antibodies (Dako) anti-mouse, anti-goat and anti-rabbit conjugated with HRP added at a dilution of 1/10,000. Equal loading of protein content was verified using alpha-tubulin as a housekeeping protein and bands visualised using ECL detection system (GE Healthcare, UK). Bands were measured using Image J densitometry, and normalised to those of loading control (alpha tubulin).

### High Resolution Respirometry of permeabilised muscle fibres by Oroboros

Respirometry studies in tibialis anterior (TA) and soleus (SOL) skeletal muscle myofibres were performed using high-resolution respirometry (Oroboros Instruments, Innsbruck, Austria). Fresh skeletal muscle (5mg) was stored in ice-cold 2mls BIOPS buffer. Any connective tissue and adipose was discarded and 20μl of Saponin stock solution for permeabilisation of muscle fibres (5mg Saponin per ml BIOPS buffer) added. The samples were then incubated for 30 minutes at 4°C. Muscles were washed in Mir05 for 10 minutes at 4°C before drying on filter paper. Approximately 3mgs of muscle tissue was introduced into the chamber and left for >10 minutes to give a stable baseline signal, re-oxygenation occurred once O_2_ concentration reached 250μM. Sequential addition of inhibitors and substrates occurred as per Oroboros Instruments specified protocol for fatty acid oxidation.

### NAD^+^ Quantification by Colorimetric Assay

NAD^+^ was extracted from flash frozen skeletal muscle and NAD^+^/NADH quantified using the NAD^+^/NADH Assay kit (BioAssay Systems) according to the manufacturers instructions.

### Unbiased Metabolomic measurement

20mg of skeletal muscle previously flash frozen tissue from TA) or Sol was placed in dry ice. Metabolites were extracted into 0.5ml of ice-cold LCMS grade methanol:acetonitrile:water (50:30:20). Samples were pulverised and stored at 4°C for 10 minutes before centrifugation at 0°C 1300rpm for 15 minutes. Supernatant was then transferred to HPLC vials ready for LC/MS analysis. To preclude variables introduces from preparing samples separately all samples were prepared simultaneously in a randomised order and re-randomised before injection in the LC/MS.

### Biocrates Targeted Metabolite kit – Acylcarnitine measurement

Skeletal muscle (Quadriceps), liver and plasma were collected from WT and H6PDKO and WT littermates. Frozen tissue was homogenised in tubes containing ceramic beads of 1.4mm diameter using extraction solvents. The AbsoluteIDQ p150 kit (Biocrates Life Science AG, Austria) was prepared according to the manufacturers instructions. Briefly, 10μl of tissue extract being used for PITC-derivatisation of amino acids, extraction with organic solvent and liquid handling steps. Acylcarnitines were quantified with a reference to appropriate internal standards using FIA-MS/MS on a 400 QTRAP instrument (AB Sciex, Germany) coupled to a HPLC Prominence (Shimadzu Deutschland GmbH, Germany).

### Metabolomic set enrichment analysis

We conducted metabolite set enrichment analysis using MetaboAnalyst^18,19^. Significantly dysregulated metabolites from the unbiased measurement in H6PDKO compared to WT TA muscle were uploaded to Metaboanalyst. These were processed with the over-representation analysis (ORA) algorithm and the metabolic pathway associated metabolite library set. This examines the group of metabolites using Fisher’s exact test, calculating the probability of seeing a particular number of metabolites containing the biological pathway of interest in the list of metabolites.

### Statistical Analysis

Students T-test or ANOVA statistical comparisons were used with the Graphpad Software Inc. Prism version 5. Data presented as mean ±SEM with statistical significance determined as *. p<0.05, **. p<0.01, ***. p<0.001. Unpaired T-Test compared treatments or genotypes. Statistical analysis derived from real-time PCR data was determined using dCt values throughout.

## Results

### Alterations in nicotinamide metabolism in H6PDKO muscle

Global and skeletal-muscle specific knockout of H6PDH provokes a myopathy characterised by metabolic stress, abnormal SR structure, SR stress and activation of the unfolded protein response. We subjected Tibialis Anterior (TA) skeletal muscle from WT and H6PDKO mice to an unbiased metabolomic screen performed by LC/MS to better understand the role H6PD plays in muscle cell metabolism and SR redox maintenance. Those metabolites significantly dysregulated in H6PDKO TA were subject to an overrepresentation analysis algorithm comparing probability of occurrence against pathway associated libraries using MetaboAnalyst^18,20^. The most over represented pathways were those involved in nicotinamide and pyrimidine metabolism (**Figure 1A-B**). Nicotinamide (NAM) generation is essential to the salvage and biosynthesis of both NAD^+^ and NADH. Therefore, changes in NAM metabolism may indicate shifts in skeletal muscle NAD^+^ availability as a result of perturbed SR NAD(P)(H) due to H6PDH deficiency. In support of this specific metabolite changes with potential relevance to SR redox include Elevated trimethyl-L-Lysine a methylated derivative of the amino acid lysine, a constituent of nuclear histone proteins and a precursor to carnitine and fatty acid oxidation in the mitochondria. Elevated 3-Hydroxysebasicacid associated with peroxisomal disorders and glycogen storage disease^21,22^, signalling a defect in fatty acids synthesis and metabolic defects such as medium chain acyl-CoA dehydrogenase deficiency. And elevated Cis-5-Tetradecenoylcarnitine-a marker of altered beta-oxidation contributing to skeletal myopathy^23^.

**Figure 1.**
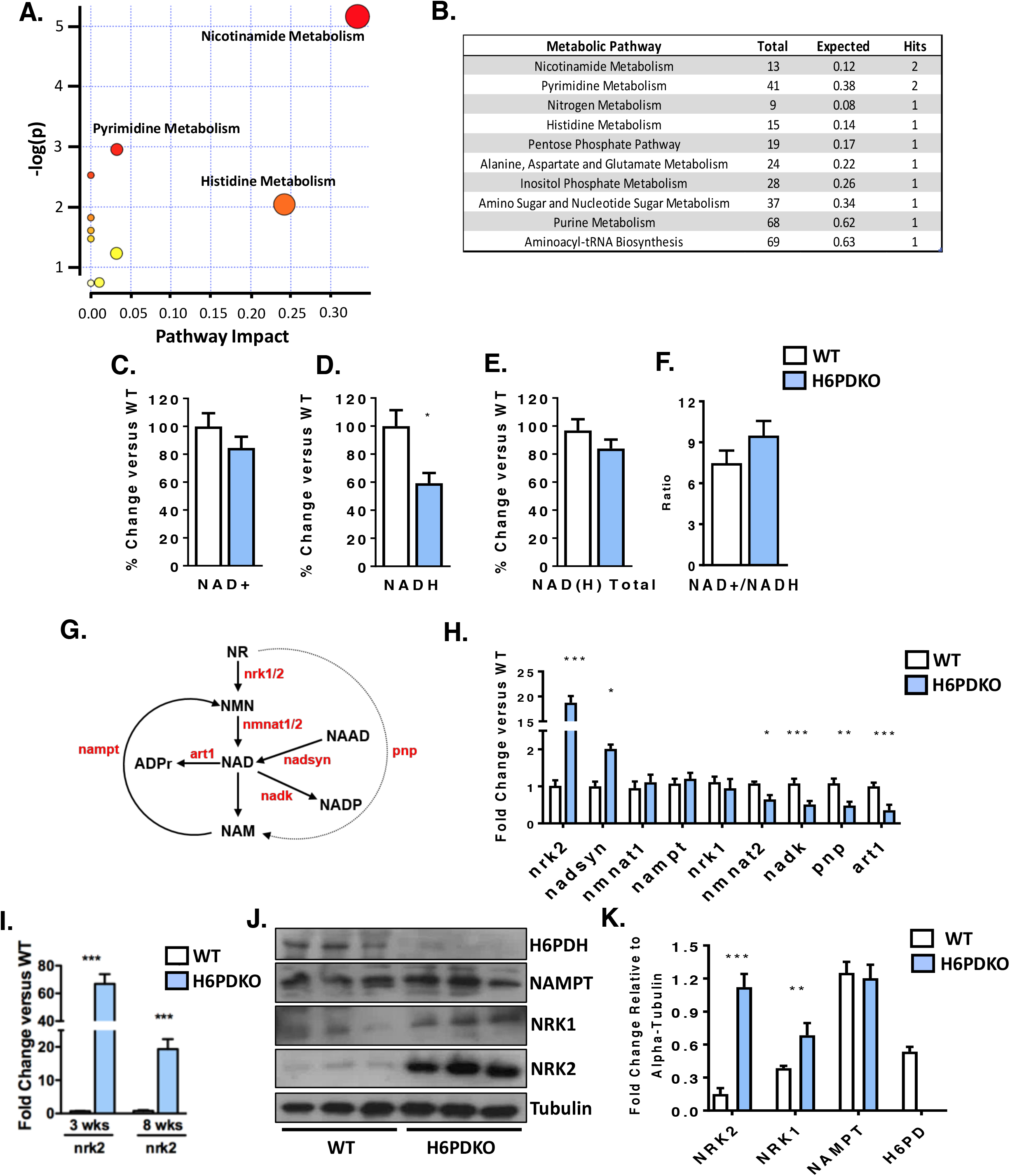
Altered NAD+ salvage pathway in H6PDKO muscle. (A) Pathways analysis of unbiased metabolomics from WT (n=3) and H6PDKO (n=3) Tibialis Anterior Skeletal Muscle. (B) Numerical output from pathway analysis of WT and H6PDH KO Tibialis Anterior Skeletal Muscle. Values shown are original value from pathway analysis. (C-F) Quantification of NAD^+^, NADH and total NAD(H) in WT (n=7) and H6PDKO (n=6) TA muscle. (G) Schematic representation of the biosynthetic generation of NAD from nicotinamide Riboside (NR) and nicotinamide (NAM) salvage, enzymes shown in red. (H) qRT-PCR of NAD^+^ synthesis and salvage genes in WT (n=7) and H6PDKO (n=7) TA. (I) Expression of NRK2 transcript in TA muscle of WT and H6PDKO mice at 3 and 8 weeks of age. (J+K) Immunoblots and quantification of WT (n=9) and H6PDKO (n=9) TA lysates. *P<0.05, **P<0.01 and ***P<0.001.

We measured NAD^+^/NADH levels and while no differences were noted in NAD^+^ levels, there was a significant 40% reduction in NADH in H6PDKO mice (**Figure 1C-F**). Critical to cellular NAD^+^ maintenance are the multiple salvage and biosynthetic enzymes culminating in NAD^+^/NADH formation (Schematic **Figure 1G**). Real-time PCR expression analysis of the genes responsible for muscle NAD^+^ biosynthesis and salvage demonstrates changes in both biosynthetic and NAD^+^ phosphorylating genes (**Figure 1H**). The most prominent adaptation was an increase in the skeletal muscle restricted *nicotinamide riboside kinase 2 (NRK2)* gene whilst the constitutively expressed salvage enzymes *NRK1* and *NAMPT* are unchanged. Responsible for the phosphorylation of the NAD^+^ precursor nicotinamide riboside into nicotinamide mononucleotide, NRK2 has previously been shown to be elevated in models of muscle energy stress and cardiomyopathy^24^. Downregulation of NAD kinase limits generation of NADP^+^, and may indicate a response to prevent NADP^+^ build up and preserve NAD for NADH production. Purine Nucleoside Phosphorylase (PNP) (converts NR to NAM) and the NAD^+^ utilising ADP-ribosyltransferase (ART1) are both downregulated, again a possible attempt to retain NAD for NADH generation. We further evaluated the expression of NAD^+^ salvage genes prior to phenotypic presentation of myopathy in 3 week old mice. At this age NRK2 was the only changed transcript, being upregulated >20-fold, suggesting this is a primary adaptive metabolic response to H6PDH deficiency (**Figure 1 I**). Western blotting confirmed elevation of NRK2 at the protein level and interestingly also suggested up regulation of NRK1 protein also (**Figure 1 J-K**). Protein expression of the rate-limiting NAMPT pathway for NAD^+^ salvage remained unchanged (**Figure 1 J-K**).

### H6PDKO skeletal muscle has reduced mitochondrial fatty acid oxidative capacity and widespread changes in acylcarnitines

Changes in NAD^+^/NADH turnover and availability can impact mitochondrial function^25–27^. Permeabilised skeletal muscle fibres from H6PDKO TA and Sol muscle were subjected to high-resolution mitochondrial respirometry. Both TA and Sol muscle have impaired oxygen consumption when exposed to palmitate as an energetic substrate indicating decreased ability to utilise substrates for fatty acid beta-oxidation and overall respiratory capacity (**Figure 2A,B**). This defect was more apparent in Sol muscle, likely representing its great mitochondrial density (**Figure 2B**). To understand if these measurements were a result of mitochondrial abundance we examined mtDNA and mitochondrial respiratory complex subunit abundance in WT and H6PDKO TA. We show no change in mitochondrial number or abundance of complexes in H6PDKO muscle to explain the defects in respiratory capacity (**Figure 2C-D**).

**Figure 2.**
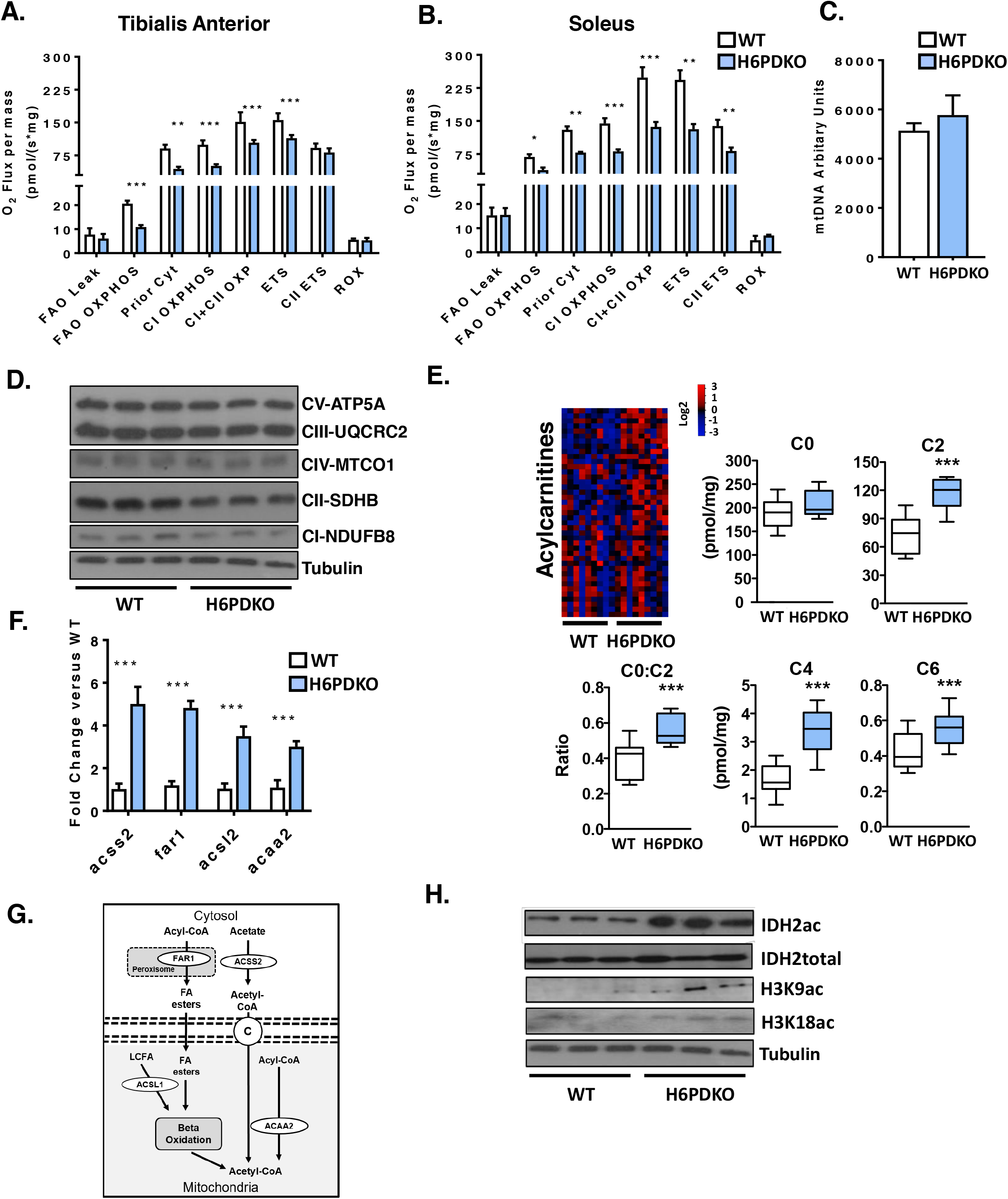
Impaired mitochondrial fatty acid oxidation in H6PDKO skeletal muscle. (A) High resolution respirometry of fatty acid oxidation in permeabilised Tibialis Anterior WT (n=3) in and H6PDKO (n=3). (B) High resolution respirometry of fatty acid oxidation using WT (n=3) and H6PDKO (n=3) permeabilised Sol. (C) Mitochondrial DNA (mtDNA) quantification of WT (n=7) and H6PDKO (n=7) muscle, measured using qRT-PCR. (D) Immunoblots WT and H6PDKO protein lysates (n=9) probed for oxidative phosphorylation enzyme subunit abundance. (E) Acylcarnitine species levels in WT (n=9) and H6PDKO (n=11) muscle measured using GC-MS/MS. Data expressed as heat-maps with log2 values representing metabolite abundance in WT and H6PDKO. Box and Whisker plots showing significantly altered short acylcarnitines. (F) qRT-PCR measurement of genes critical carnitine and fatty acids metabolism in WT (n=7) and H6PDKO (n=7) TA. (G) Schematic showing carnitine and fatty acid metabolism between cytosol and mitochondria. (H) Immunoblots of acetylated proteins within WT (n=6) and H6PDKO (n=6) skeletal muscle. *P<0.05, **P<0.01 and ***P<0.001.

To further examine the basis for increased reliance on lipid burning and in the face of impaired mitochondrial fatty acid oxidation we conducted a targeted lipidomic analysis of serum, liver and quadriceps from H6PDKO and WT mice (**Supplementary Figure 1**). Liver and serum from H6PDHKO showed no significant changes in lipid profile. However, a striking effect for elevated short-chain acylcarnitine (C2, C3, C4, C5, C6 & C10:1) abundance in H6PDKO muscle tissue was seen. While free-carnitine was not significantly different, the ratio of free carnitine (C2:C0) was increased, possibly signifying increased production or block in use of acetyl Co-A in H6PDKO muscle. (**Figure 2E**).

Using whole genome expression array data, validated with real-time PCR, we show that H6PDKO muscle have significant elevations in genes transcripts ACSS2, FAR1, ACSL1, and ACAA2, which converge on acylcarnitine and acetyl-CoA metabolism (**Figure 2F, G**)^12^. Acetyl-CoA produced during fatty acid oxidation is important to the levels of histone acetylation and the regulation of gene expression^28,29^. Accumulation of mitochondrial acetyl Co-A can lead to non-enzymatic acetylation of multiple proteins associated with energy metabolism^30,31^. Protein lysates collected from H6PDKO muscle show increases in markers of increased mitochondrial-Isocitrate Dehydrogenase 2 (IDH2) and nuclear histone 3 lysine 9 and lysine 18 acetylation, endorsing the fatty acid oxidation and acyl-CoA changes observed in gene expression, mitochondrial and metabolomic measures (**Figure 2H**).

### Rescuing NAD^+^ metabolism using nicotinamide riboside

We reasoned that increased drive towards Acetyl-CoA production and acylcarnitine accumulation leads to protein acetylation and inhibition of mitochondrial function^32^. Increasing NAD^+^ availability for the mitochondrial NAD-dependent protein decetylases, such as SIRT3, could relieve the burden of protein acetylation to improve energy metabolism^33–36^. To test this we took advantage of the elevated expression of the NRK pathway and supplemented mice with NR (IP, 400mg/kg twice daily over 5 consecutive days), a naturally occurring B3 vitamin NAD^+^ precursor. Quantification of NAD(H) levels in the skeletal muscle of mice receiving NR showed it to enhance total NAD(H), rescuing NADH levels (**Figure 3 A-D**). Furthermore, the ability to elevate NAD(H) was more pronounced in H6PDKO muscle with upregulated NRK pathway activity.

**Figure 3.**
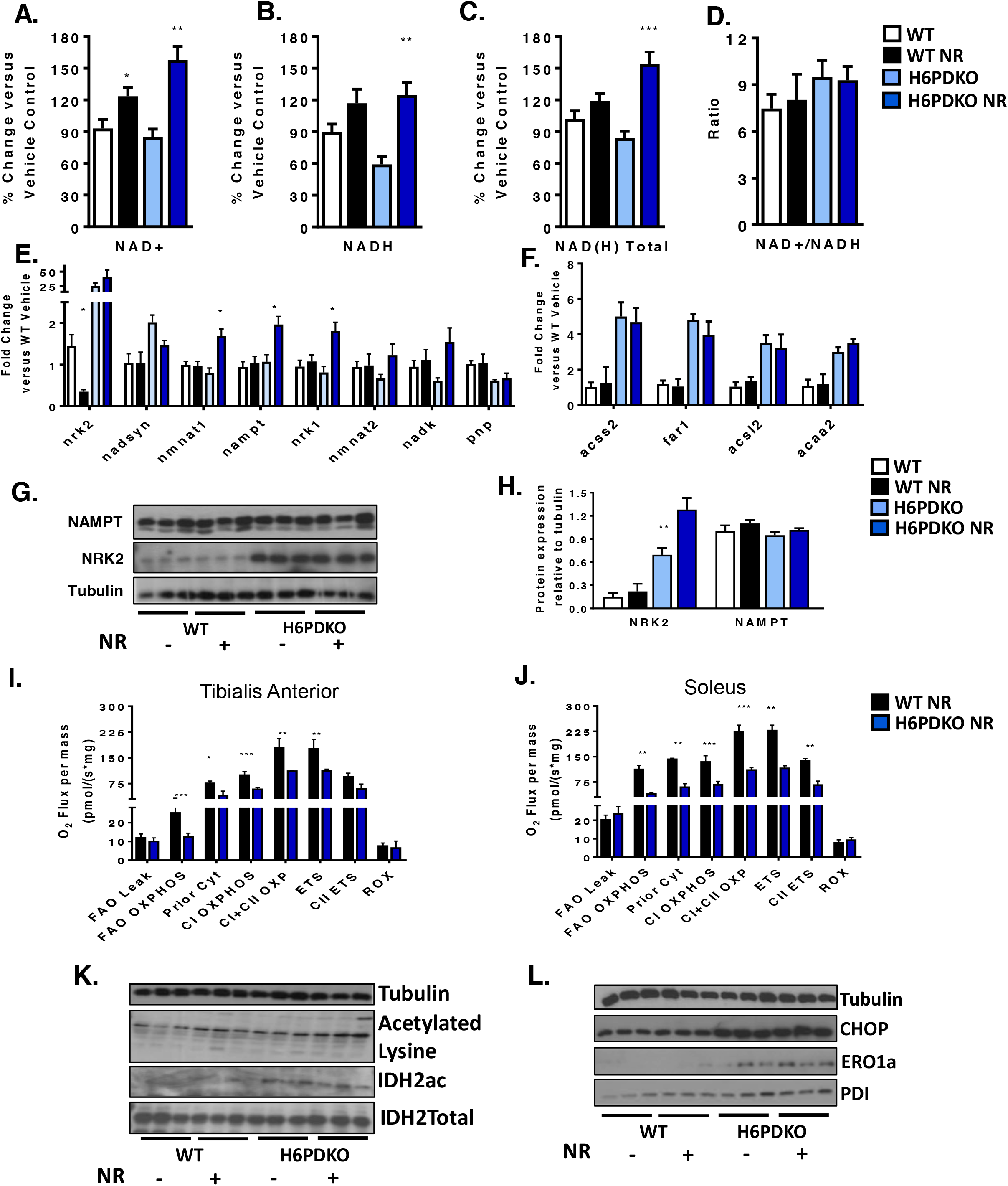
NR supplementation, NAD+ salvage and mitochondrial function in H6PDKO muscle. (A-D) NAD^+^, NAD(H) and total NAD(H) quantification in skeletal muscle of WT and H6PDKO +/− intraperitoneal (i.p.) Nicotinamide Riboside (NR) (n=6-9). (E) qRT-PCR data of genes involved in the biosynthesis of NAD^+^ and salvage of NAM in WT and H6PDKO muscle +/− i.p. NR (n=6-9). (F) qRT-PCR data of genes critical carnitine and fatty acids metabolism in WT and H6PDKO muscle +/− i.p. NR (n=6-9). (G-H) Immunoblots and quantification of the NAM salvage protein NAMPT and the skeletal muscle specific protein NRK2 (n=6). (I) High resolution respirometry of fatty acid oxidation in permeabilised Tibialis Anterior WT in and H6PDKO i.p NR (n=3). (I) High resolution respiration for fatty acid oxidation using WT and H6PDKO after i.p NR permeabilised Sol (n=3). (J) Immunoblots showing ER stress regulator CHOP and protein folding factors PDI and ERO1a in WT and H6PDKO +/− i.p NR. (K) Immunoblots showing total lysine acetylation and IDH acetylation in WT and H6PDKO muscle +/− i.p NR. (n=6-9). *P<0.05, **P<0.01 and ***P<0.001.

NR had an effect to decrease NRK2 expression in WT but not H6PDKO muscle. However, NR did lead to increased expression of NMNAT1, NAMPT and NRK1 in H6PDKO muscle, which was not seen in WT (**Figure 3E**). NAD^+^ boosting has no impact to suppress Acyl-CoA and lipid metabolism gene expression in H6PDKO mice (**Figure 3F**). Protein quantification of NAMPT and NRK2 showed no change in abundance (**Figure 3G+H**). Following NR supplementation we again performed high resolution mitochondrial respirometry of fatty acid oxidation and found no improvement in mitochondrial O_2_ flux in NR supplemented WT or H6PDKO muscle (**Figure 3 I-J**).

To further assess the effects of augmented NAD(H) availability we examined protein acetylation in response to NR. Global and IDH2 acetylation were maintained at untreated H6PDKO levels (**Figure 3L**). Finally, H6PDKO muscle has extensive activation of the unfolded protein response, potentially a consequence of altered NADPH/NADP^+^ ratio and impaired protein folding^13,37,38^. UPR markers CHOP and the protein folding regulators ERO1a and PDI levels were evaluated and shown to remain elevated in H6PDKO mice despite NR supplementation and increased NAD(H) availability (**Figure 3K**). These data imply that perturbed SR NAD(P)(H) as a function of H6PDH deficiency cannot be overcome by boosting NAD(H) availability according to this study design.

### NRK2 is dispensable in H6PDKO myopathy

We now investigated the importance of the NRK2 pathway in defending muscle function in the absence of H6PDH using double knockout (DKO) mice. We reasoned DKO mice would have an exacerbated myopathy, and may even reduce survival. However, DKO mice were viable, survived at the anticipated frequency and generally indistinguishable from H6PDKO mice. Quantification of NAD^+^ levels revealed that DKO TA has significantly decreased NAD^+^ levels compared to WT and to a greater degree than in H6PDH deficiency alone, with NADH and total NAD(H) remaining depressed at H6PDKO levels (**Figure 4A-D**). The increased expression of NRK2 would therefore appear to be able to defend aspects of the NAD metabolome in H6PDH deficiency. NRK2 deficiency alone had minimal impact in WT mice as previously reported, and likely due to compensatory activity of the NRK1 enzyme^39^. DKO skeletal muscle exhibited myopathy and atrophy to an equivalent degree as in H6PDKO (**Figure 4 E-G**). Expression of genes for NAD^+^ biosynthesis and Acyl-CoA metabolism revealed no significant differences in the DKO muscle compared to H6PDKO, with NRK2KO displaying no change compared to WT (**Figure 4H, I**).

**Figure 4.**
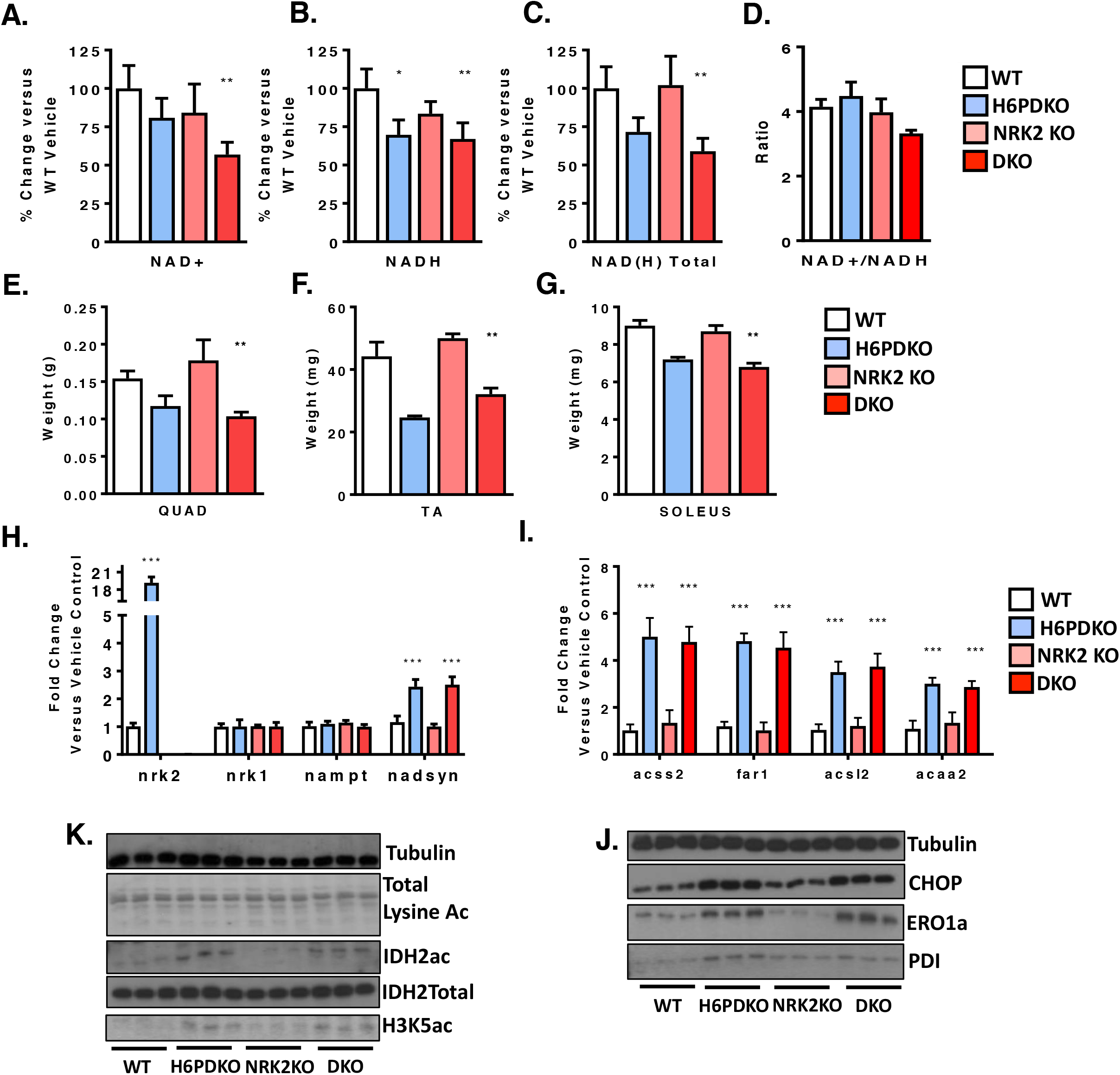
Molecular and phenotypic analysis of H6PDH-NRK2 Double knockout mice. (A-D) NAD^+^, NADH and total NAD(H) levels in WT, H6PDKO, NRK2KO and DKO (H6PDKO-NRK2KO) TA muscle (n=3-6). (E-G) Skeletal muscle tissue weights from WT, H6PDKO, NRK2KO and H6-NRK2 Double Knockout (DKO) (n=3-6). (H) qRT-PCR of NAD^+^ biosynthetic gene expression in WT, H6PDKO, NRK2 KO and DKO TA (n=3-6). (I) qRT-PCR of mitochondrial and Acyl-CoA genes in TA of WT, H6PDKO, NRK2 KO and DKO (n=3-6). (J) qRT-PCR of ER/SR stress response genes in WT, H6PDKO, NRK2KO and DKO TA (n=3-6). (F-K) Immunoblots of CHOP and protein folding factors PDI and ERO1a in WT, H6PDKO, NRK2 KO and DKO muscle lysates (n=3-6). (L) Immunoblots of total lysine acetylation, IDH2 acetylation in WT, H6PDKO, NRK2KO and DKO muscle protein lysates (n=3-6). *P<0.05, **P<0.01 and ***P<0.001.

Finally we examined protein acetylation and ER stress and UPR markers, all of which were, again, unchanged from the levels seen in H6PDKO muscle (**Figure 4J+K**).

## Discussion

In this work we show that upregulation of NRK2 mediated salvage of NR into NAD^+^ is an early adaptation to perturbed muscle SR NAD(P)(H) homeostasis and impaired mitochondrial energy production in H6PDH deficiency. This study demonstrates that NR supplementation can defend NAD(H) levels within H6PD deficient muscle, but this, and ablation of the stress responsive NRK2 pathway have little impact to limit or worsen myopathy.

The nicotinamide riboside kinases 1 and 2 (NRKs1/2) form part of the salvage pathways that convert NR into NMN through to NAD^+^^39^. Importantly, extracellular NMN conversion to NAD^+^ is NRK dependent, requiring it to be dephosphorylated to NR to be utilised as an NAD^+^ precursor^40^. NRK1 is thought to be the dominant and rate-limiting source of NR phosphorylation^39,40^ in most tissues, with NRK2 expression being limited to skeletal and cardiac muscle tissues^37^. Upregulation of NRK2 has been observed as a primary response to metabolic energy stress associated with depletion of NAD^+^ in genetic models of cardiomyopathy and in a model of post-injury skeletal muscle repair^24,39,41^. Despite its limited expression in non-muscle tissues, NRK2 has also been shown to be induced in axonal neurons in response to damage, further supporting a role in response to cellular damage or metabolic stress^42^.

Supplementation of the vitamin B3 NR as an NAD^+^ precursor is well tolerated in both mice and humans and effective in elevating muscle NAD^+^ levels and alleviating aspects of the pathology seen in muscular dystrophy, mitochondrial myopathy and ageing^43–45^. Indeed, NR supplementation has proven to be effective in alleviating cardiac defects associated with genetic cardiomyopathies’^24,41^. We report that NR was successful at enhancing NAD(H) levels in H6PDKO muscle but unable to rescue physiological or molecular defects associated with the metabolic myopathy, despite elevated NRK2 activity. It is reasonable to suggest more sustained NR delivery, beyond the 5 day protocol used here, may have revealed effects.

We also show that abolishing NRK2 upregulation in H6PDKO muscle confers no discernible disadvantage, with DKO mice indistinguishable from H6PDKO mice, despite further deterioration in NAD+ availability. We have previously shown that NRK1 is able to compensate for nrk2 deficiency in terms of NAD^+^ salvage, and as NRK1 protein is also upregulated in H6PDKO muscle, may again do so in this context^39^. Also, muscle-specific NAMPT KO mice are viable and only develop myopathy after several months despite NAD^+^ levels at 90-95% those of WT mice^27^. We were unable to ascertain the levels of NR available to the NRK2 pathway in H6PD deficiency, and suggest that despite NRK2 elevation, NR availability will be limiting, even with supplementation^46^.

Upregulation of NRK2 may be as an early adaptive response to metabolic stress and the need to defend NAD^+^ availability. In cardiomyocytes the Nrk2 gene is activated by energy stress and NAD^+^ depletion through AMP-activated protein kinase and peroxisome proliferator-activated receptor α dependant mechanisms^24,47^. This may be important in exercise adaptation as aged NRK2 KO mice show maladaptive metabolic responses to exercise in cardiac and skeletal muscle^46^.

The role NRK2 may play as a direct response to perturbed SR NAD(P)(H) homeostasis remains less clear given NAD^+^ precursor supplementation had no impact over acetlycarnitine metabolism or mitochondrial function. It is possible that elevated NRK2 is direct response to perturbed SR NAD(P)(H) homeostasis and regulated through a mechanism distinct from that initiated upon energy stress and NAD^+^ depletion. This notion may support the concept of SR sensing and exchange of nucleotides, nucleosides or other precursors involved in NAD(P)(H) homeostasis. Recent findings have identified transporters able to carry NAD^+^ and NMN across biological membranes. Importantly, NAD^+^ has been shown to be transported into the mitochondria and as such increases the potential for other compartments, such as the ER/SR, to transport NAD^+^ and associated precursors^48^. Indeed, Slc12a8 has recently been identified as an NMN transporter regulating intestinal NAD^+^ metabolism^49^. Although an NR transporter has been identified in yeast, there remains no conclusive evidence for a dedicated NR transporter in mammalian cells^50^.

## Conclusion

These data identify activation of the NRK2 pathway, and concurrent changes in NAD^+^ metabolism as an early response to perturbations in ER/SR NAD(P)(H) homeostasis as a result of H6PDH deficiency in skeletal muscle. Whether upregulation of NRK2 mediated NAD(P)(H) salvage is a response to mitochondrial dysfunction and energy stress, or directly to perturbation in NADP(H) availability remains to be determined. While H6PDH deficiency results in a deteriorating and complex metabolic and structural myopathy that cannot be rescued with NAD^+^ supplementation, it remains an intriguing model linking muscle SR NAD(P)(H) mediated redox and energy metabolism.

## List of Abbreviations

ER/SR: : Endoplasmic/Sarcoplasmic Reticulum
H6PD: : Hexose-6-Phosphate Dehydrogenase
NAD^+^: : Nicotinamide Adenine Dinucleotide
NMN: : Nicotinamide Mononucleotide
NR: : Nicotinamide Riboside
NRK: : Nicotinamide Riboside Kinase
TA: : Tibialis Anterior
SOL: : Soleus

## Author contributions

CLD, AEZ and GGL conceived the study. CLD, GGL and DAT wrote the manuscript. Experimental work was performed by CLD, AEZ, LOB, RSF, YH, DC, AG, DW, JA.

## Acknowledgements

We thank Chromadex (Irvine, California) for providing nicotinamide riboside.

## Availability of data and materials

Datasets used in this study are available from the author upon request.

---

**Supplementary Figure 1.**
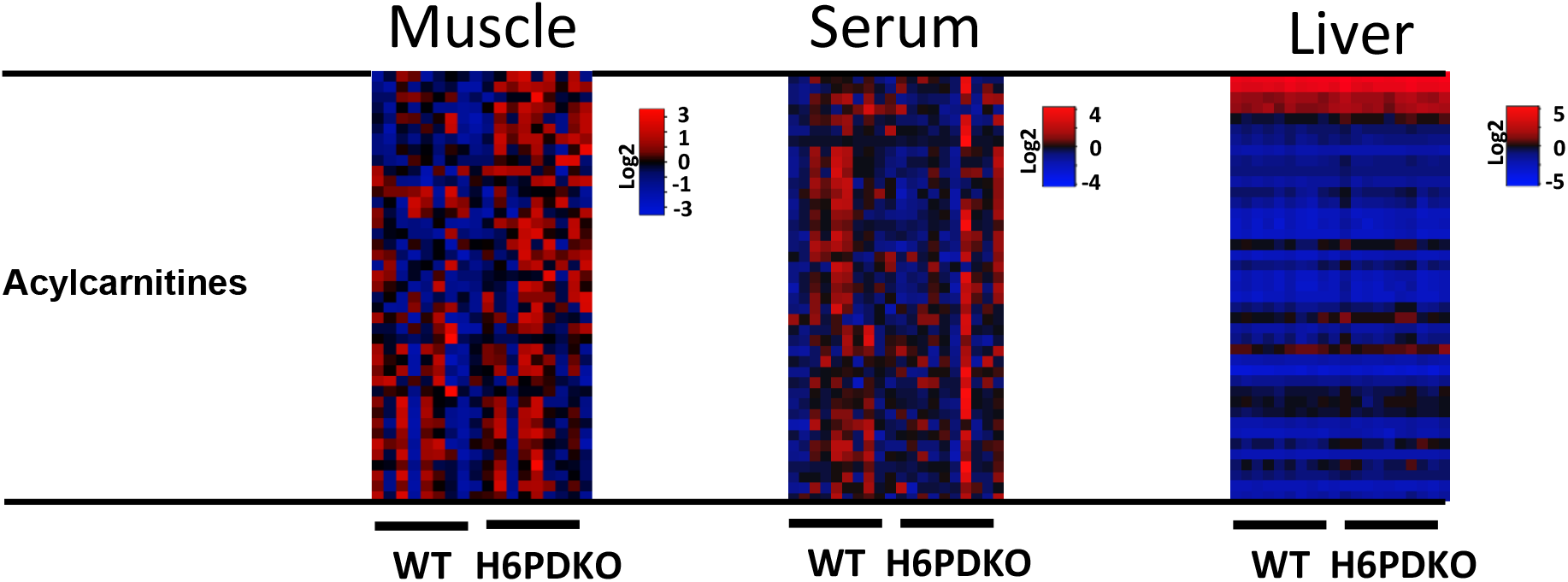
Acylcarnitine measurment of WT and H6PDKO skeletal muscle, serum and liver.

